# Chronic Interleukin-1 exposure triggers selective expansion of *Cebpa*-deficient multipotent hematopoietic progenitors

**DOI:** 10.1101/2020.03.25.008250

**Authors:** Kelly C. Higa, Andrew Goodspeed, James S. Chavez, Vadym Zaberezhnyy, Jennifer L. Rabe, Daniel G. Tenen, Eric M. Pietras, James DeGregori

**Author notes:** Correspondence to Eric Pietras and James DeGregori.

## Abstract

The early events that drive hematologic disorders like clonal hematopoiesis, myelodysplastic syndrome, myeloproliferative neoplasm, and acute myeloid leukemia are not well understood. Most studies focus on the cell-intrinsic genetic changes that occur in these disorders and how they impact cell fate decisions. We consider how chronic exposure to the pro-inflammatory cytokine, interleukin-1β (IL-1β), impacts *Cebpa*-deficient hematopoietic stem and progenitor cells (HSPC) in competitive settings. We surprisingly found that *Cebpa*-deficient HSPC did not have a hematopoietic cell intrinsic competitive advantage; rather chronic IL-1β exposure engendered potent selection for *Cebpa* loss. Chronic IL-1β augments myeloid lineage output by activating differentiation and repressing stem cell gene expression programs in a *Cebpa*-dependent manner. As a result, *Cebpa*-deficient HSPC are resistant to the pro-differentiative effects of chronic IL-1β, and competitively expand. These findings have important implications for the earliest events that drive hematologic disorders, suggesting that chronic inflammation could be an important driver of leukemogenesis and a potential target for intervention.

**Summary:** Higa *et al*. show that chronic interleukin-1β exposure primes hematopoietic stem and progenitor cells for myelopoiesis by upregulating myeloid differentiation programs and repressing stem gene programs in a *Cebpa*-dependent manner. Consequently, interleukin-1 potently selects for *Cebpa* loss in hematopoietic stem and progenitor cells.

## Introduction

Hematologic disorders such as clonal hematopoiesis (CH), myelodysplastic syndrome (MDS), myeloproliferative neoplasm (MPN), and acute myeloid leukemia (AML) are thought to originate from multipotent hematopoietic stem and progenitor cells (HSPC), which are responsible for life-long regeneration of the blood system. Understanding the earliest events that drive these disorders is thus important to understand the etiology of the malignancy and to devise potential intervention strategies.

Risk factors associated with hematologic disorders include age and prior exposure to radiation or chemotherapy, conditions characterized by underlying inflammation (Bowman et al., 2018). Prior studies have shown that aging, radiation and chronic inflammation promote HSPC differentiation at the expense of self-renewal (Fleenor et al., 2015a; Mejia-Ramirez and Florian, 2020; Pietras, 2017). The pro-inflammatory cytokine, interleukin-1 (IL-1) is important in regulating hematopoiesis and the immune response in the acute setting but can drive pathology in the chronic setting (Mantovani et al., 2019). For instance, IL-1 can promote AML expansion in *ex vivo* culture (Carey et al., 2017), and increased expression of IL-1 receptor accessory protein is associated with poor prognosis in AML (Barreyro et al., 2012).

Many oncogenic mutations associated with AML are known to block differentiation and/or enhance self-renewal (Cancer Genome Atlas Research et al., 2013; Corces et al., 2017; Kelly and Gilliland, 2002). *CEBPA* is frequently altered in MDS/MPN and AML by loss-of-function (LOF) mutations or by (epi)genetic alterations that suppress *CEBPA* expression, together accounting for ∼70% of molecular alterations in *de novo* AML (Fig. 1A, (Cancer Genome Atlas Research et al., 2013; Ernst et al., 2010; Gu et al., 2018; Guo et al., 2012; Pabst et al., 2001a; Perrotti et al., 2002; Zheng et al., 2004)). Further, *Cebpa* LOF leads to a block in myeloid differentiation, contributing to AML-like diseases in mouse models (Bereshchenko et al., 2009; Kirstetter et al., 2008; Pabst et al., 2001b; Porse et al., 2005; Zhang et al., 1997; Zhang et al., 2004). Therefore, *CEBPA* LOF represents a common mechanism of hematopoietic dysregulation across hematologic disorders. Mechanism(s) driving selection for the *CEBPA* LOF phenotype in hematologic disorders are poorly understood.

**Figure 1:**
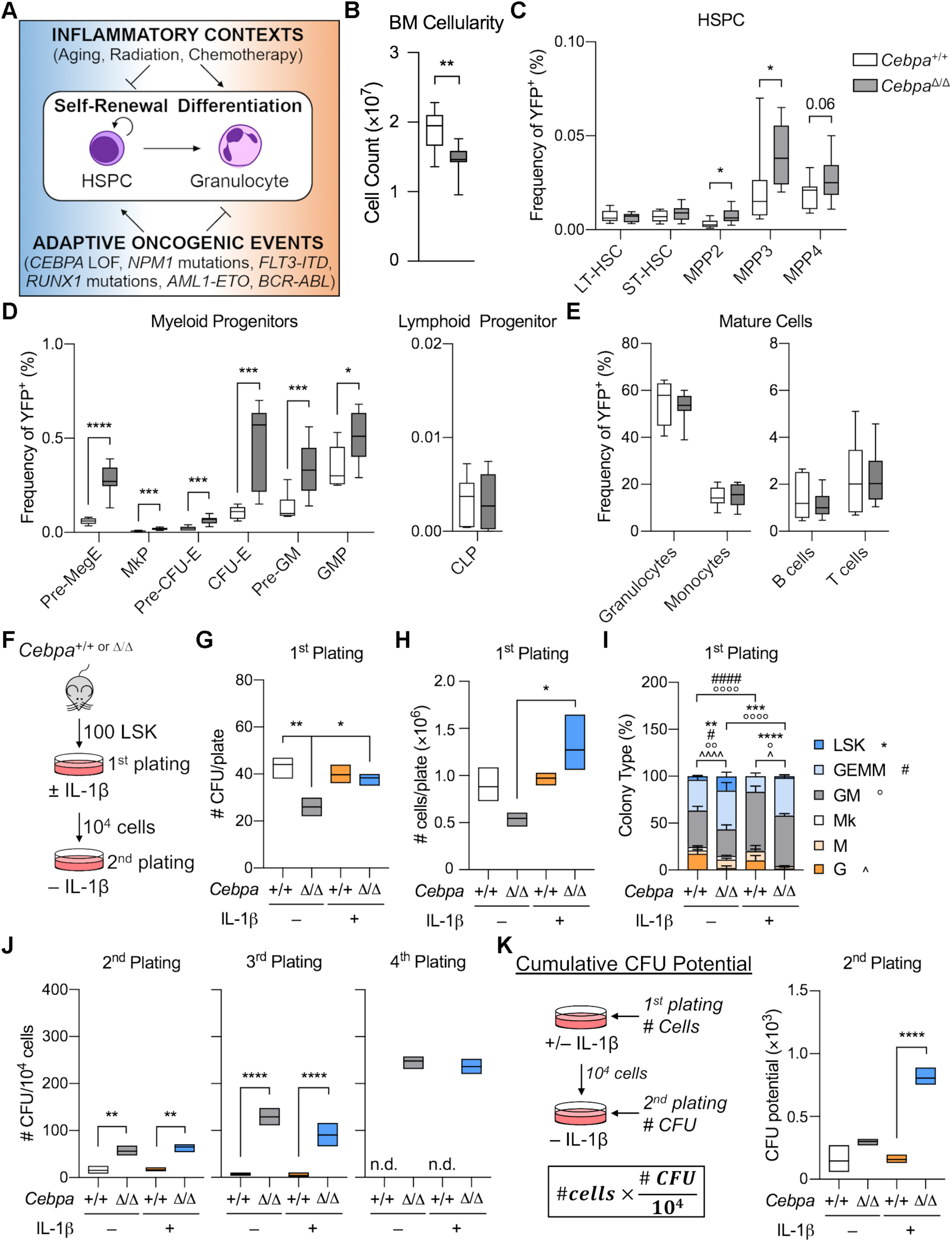
HSPC-specific *Cebpa* deficiency enhances MPP activity. A) Model of inflammation-driven HSPC differentiation that selects for phenotypes that prevent HSPC differentiation and/or promote self-renewal (oncogenic mutations indicated have all been shown to impair *Cebpa* expression (Cancer Genome Atlas Research et al., 2013; Ernst et al., 2010; Gu et al., 2018; Guo et al., 2012; Pabst et al., 2001a; Perrotti et al., 2002; Zheng et al., 2004)). B-E) Hematopoietic characterization of *Cebpa*^+/+^ and *Cebpa*^Δ/Δ^ mice (n=9 per genotype) 7 days post-deletion: B) Bone marrow cellularity from 1 femur and 1 tibia per mouse; C) HSPC frequency; D) Committed progenitor frequency; E) Mature cell frequency. F) Experimental design for LSK CFU assay. G-K) LSK CFU assay data (representative of 3 separate experiments with n=3 per group): G) CFU counts per plate for initial plating; H) Total cell counts per plate for initial plating; I) Frequencies of CFU types for initial plating; J) CFU counts per plate on re-plating 10^4^ cells (n.d.: not done because too few cells were recovered); K) Calculation and data for Cumulative CFU potential for 2^nd^ plating. All data are presented as mean ± s.d. B-E) Unpaired Mann-Whitney u-test. G-K) Two-way ANOVA with Tukey’s multiple comparisons test. *, p<0.05; **, p<0.01; ***, p<0.001; ****, p<0.0001, except in I where symbols for CFU types are indicated. Could not perform statistics for 4^th^ replating due to missing data points.

Analysis of steady-state hematopoiesis suggests that multipotent progenitors (MPP) derived from hematopoietic stem cells (HSC) are major daily contributors to blood production, and are long-lived under unperturbed conditions in mice (Busch et al., 2015; Sawen et al., 2016; Sun et al., 2014). Thus, in order to understand the origin of hematologic disorders, it is important to consider both HSC and MPP, collectively referred to as HSPC, as potential oncogenic reservoirs, and to consider how they interact with their microenvironment.

We hypothesized that chronic inflammation may drive selection for adaptive phenotypes like *Cebpa* LOF that counteract HSPC differentiation (Fig. 1A). To test this hypothesis, we modeled the impact of chronic inflammation on competition between wild-type (WT) and *Cebpa*-deficient HSPC *in vivo*, by treating chimeric mice with IL-1β for 20 days. *Cebpa*-deficient HSPC are resistant to the pro-differentiative effects of IL-1β, thereby providing a fitness advantage in this setting, and leading to their preferential expansion in the bone marrow (BM). These data illustrate the potential for inflammation to alter the intrinsic transcriptional landscape of HSPC and to drive the selective fitness of potentially oncogenic HSPC in the BM.

## Results and Discussion

### HSPC-specific *Cebpa* deficiency causes expansion of multipotent progenitors

To understand how *Cebpa* disruption impacts HSPC fitness in a context-dependent manner, we crossed *Cebpa*^f/f^ mice (Zhang et al., 2004) with *R26R-LSL-EYFP* and *HSC-SCL-Cre-ER*^*T*^ transgenic mice (Gothert et al., 2005) to achieve traceable, tamoxifen-inducible, HSPC-specific knockout of *Cebpa* (herein referred to as *Cebpa*^Δ/Δ^). To induce *Cebpa* excision, we treated mice with 2.5 mg tamoxifen for 3 days and analyzed mice 7 days later (Fig. S1A). We confirmed that YFP accurately reports *Cebpa* excision by genotyping individual colonies formed on methylcellulose from YFP^+^ HSPC-enriched Lineage^−^Sca1^+^cKit^+^ (LSK) cells (Fig. S1B, and data not shown). Two days after tamoxifen treatment, the LSK population was highly and specifically labeled, followed by the appearance of YFP^+^ cells within more differentiated populations by day 7 (Fig. S1C).

To assess the impact of *Cebpa* deficiency on hematopoietic lineage distribution in this model, we analyzed BM and peripheral blood (PB) from *Cebpa*^+/+^ and *Cebpa*^Δ/Δ^ mice (Fig. S1D-F). BM cellularity was significantly reduced in *Cebpa*^Δ/Δ^ mice (Fig. 1B). *Cebpa* deficiency did not significantly affect the frequency of phenotypically defined long-term HSC (LT-HSC) or short-term HSC (ST-HSC), but significantly increased the frequency of megakaryocyte/erythroid (MegE)-biased MPP2 and granulocyte/monocyte (GM)-biased MPP3, with a trending increase in lymphoid-biased MPP4 (Fig. 1C). *Cebpa*^Δ/Δ^ mice also had a significant build-up of MegE-lineage progenitors and GM-lineage progenitors, suggestive of a block in myeloid lineage differentiation at this level. We did not observe differences in the common lymphoid progenitor (CLP) (Fig. 1D). *Cebpa*^Δ/Δ^ mice had no significant changes in the frequency of mature BM cell populations (Fig. 1E) or mature PB cells, except for a significant reduction in red blood cell (RBC) counts (Fig. S1F). The lack of changes in mature BM cells or in LT-HSC or ST-HSC differs from previous studies (Ye et al., 2013; Zhang et al., 2004). This is likely due to using an HSPC-specific Cre rather than *Mx1*-Cre, where the effect of a differentiation block is delayed due to the slower turnover of HSCs and MPPs. This mouse model thus corroborates previous findings that C/EBPα acts during myeloid lineage specification upstream of GMP (Zhang et al., 2004) and pre-GMs (Pundhir et al., 2018).

### HSPC-specific *Cebpa* deficiency enhances MPP colony forming potential

IL-1β promotes precocious myeloid differentiation with concomitant loss of HSC self-renewal (Pietras et al., 2016). Because *Cebpa* deficiency blocks myeloid differentiation upstream of GMP and pre-GM, we hypothesized that *Cebpa*^Δ/Δ^ LSK cells would retain serial replating activity when exposed to IL-1β. We sorted 100 LSK cells from *Cebpa*^+/+^ or *Cebpa*^Δ/Δ^ mice and performed methylcellulose colony forming unit (CFU) assays with or without IL-1β (Fig. 1F). We assessed colony types and numbers using both morphological scoring and flow cytometry (data not shown). While IL-1β did not affect the number of CFU or total cell numbers from *Cebpa*^+/+^ LSK cells on initial plating (Fig. 1G-H), it did skew output toward significantly more differentiated CFU-GM colonies (both Mac1^low^Gr1^+^ and Mac1^high^Gr1^+^ by flow cytometry) with significantly fewer CFU-GEMM (Lin^−^Sca1^−^cKit^+^ by flow cytometry) colonies (Fig. 1I), consistent with previous studies using purified HSC (Pietras et al., 2016). In the absence of IL-1β, *Cebpa*^Δ/Δ^ LSK cells formed significantly fewer CFU, producing fewer mature CFU-G (Mac1^low^Gr1^+^ by flow cytometry) than *Cebpa*^+/+^ LSK cells while retaining significantly more immature CFU-Blast (Lin^−^Sca1^+^cKit^+^ by flow cytometry) (Fig 1G-I). In contrast, IL-1β significantly increased the clonogenic potential of *Cebpa*^Δ/Δ^ LSK cells (Fig. 1G). Even in the presence of IL-1β, *Cebpa*^Δ/Δ^ LSK cells maintained significantly more multipotent CFU-GEMM and produced fewer mature CFU-GM and CFU-G than that of *Cebpa*^+/+^ (Fig. 1I).

To functionally assess HSPC clonogenic potential, we serially replated 10^4^ cells on methylcellulose without IL-1β (Fig. 1F). On the 2^nd^ plating, *Cebpa*^Δ/Δ^ cells formed significantly more CFU than *Cebpa*^+/+^ cells, regardless of prior IL-1β exposure. *Cebpa*^+/+^ cells were not maintained past the 3^rd^ replating, whereas *Cebpa*^Δ/Δ^ cells continued to expand in number up to a 4^th^ replating (Fig. 1J). These data suggest that the absence of C/EBPα prevents differentiation while preserving progenitor activity even in the context of IL-1β.

For the initial plating, IL-1β significantly increased the number of CFU and total cells produced from *Cebpa*^Δ/Δ^ LSK cells (Fig. 1G-H). For subsequent rounds of plating, *Cebpa*^Δ/Δ^ cells had similar clonogenic potential regardless of prior IL-1β exposure. In order to account for the initial expansion difference in *Cebpa*^Δ/Δ^ LSK cells with and without IL-1β, we calculated a “cumulative CFU potential” by multiplying the total number of cells from the 1^st^ plating by the CFU formed per 10^4^ cells at replating, demonstrating that IL-1β significantly increased cumulative CFU potential of *Cebpa*^Δ/Δ^ LSK cells (Fig. 1K). Together, these data suggest that IL-1β potentiates initial *Cebpa*^Δ/Δ^ HSPC expansion, which ultimately results in increased cumulative CFU potential.

### *Cebpa* deficiency confers a fitness advantage in the context of chronic IL-1β in competitive transplants

The IL-1β-enhanced colony-forming activity we observed led us to hypothesize that IL-1β may engender a fitness advantage for *Cebpa*^Δ/Δ^ LSK cells *in vivo*. To test this concept, we competitively transplanted sorted LSK cells from donor *Cebpa*^+/+^ or *Cebpa*^Δ/Δ^ mice with C57BL/6 (CD45.2) competitor bone marrow into congenic Boy/J (CD45.1) recipients, allowing us to track the donor, competitor, and recipient populations by flow cytometry (Fig. 2A-B). In order to isolate the effects of chronic IL-1β in the transplant setting, we used mild busulfan conditioning, which we previously showed to be less inflammatory and less disruptive to the hematopoietic system than irradiation (Henry et al., 2015). Because busulfan does not significantly deplete mature hematopoietic cells, we also treated recipients with CD4 and CD8 T cell depleting antibodies to prevent rejection of *Cebpa*^+/+^ or *Cebpa*^Δ/Δ^ donor cells expressing YFP, a potential foreign antigen. At 3 weeks post-transplant we found that PB chimerism ranged between 1-7% and was significantly lower for *Cebpa*^Δ/Δ^-derived cells (Fig. 2C). Thus, this system models early stages of leukemogenesis whereby rare oncogenically-initiated cells must compete with more abundant normal progenitors.

**Figure 2:**
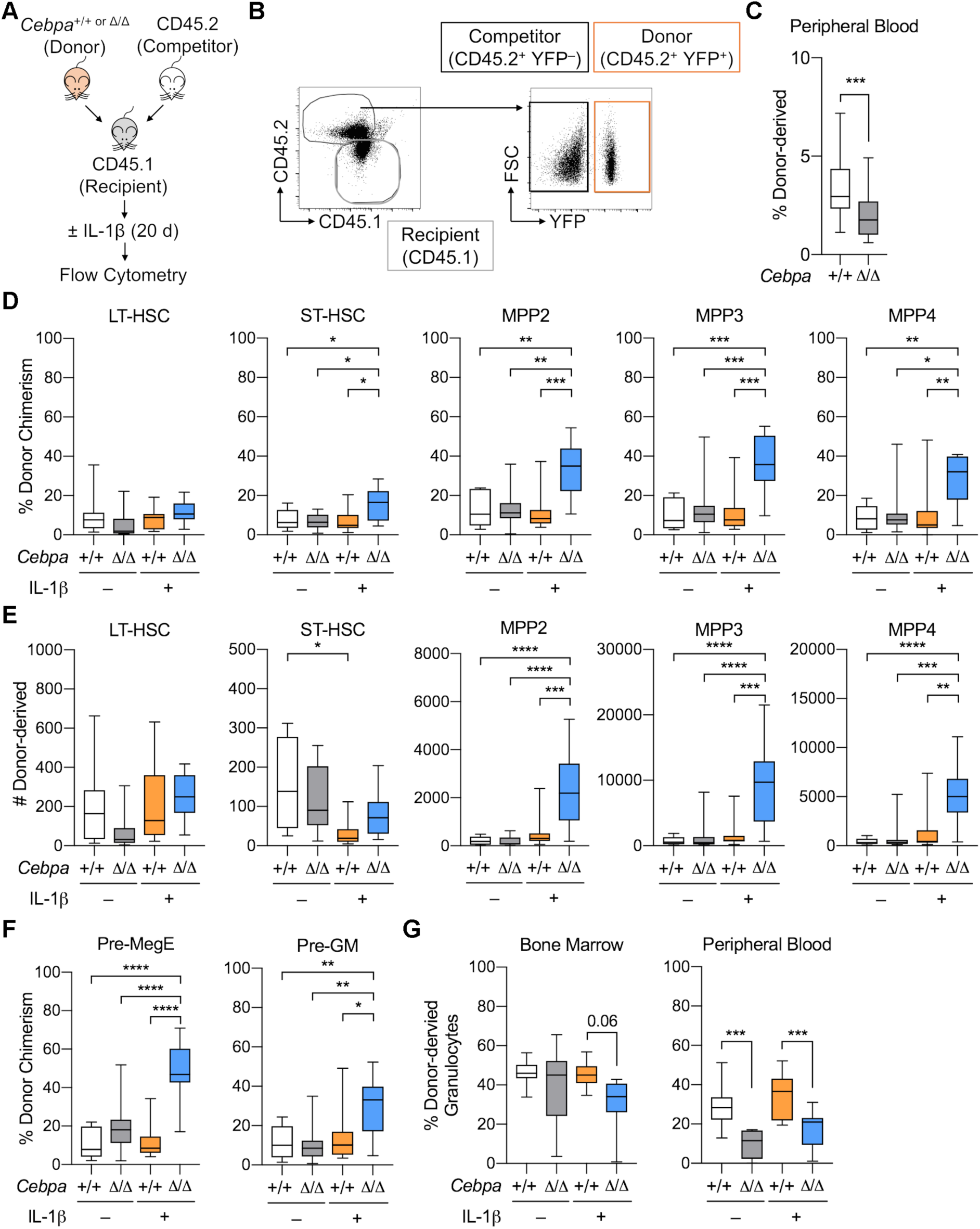
*Cebpa* deficiency confers a fitness advantage in the context of chronic IL-1β in competitive transplants. A) Experimental design for competitive bone marrow transplants. B) Representative flow cytometry plots to identify donor (CD45.2^+^YFP^+^), competitor (CD45.2^+^YFP^−^), and recipient (CD45.1^+^) populations. C) Initial engraftment in the periphery was assessed by frequency of donor-derived (CD45.2^+^YFP^+^) leukocytes at 3 weeks post-transplant. D-F) Following 20 days of treatment with vehicle or IL-1β, bone marrow was analyzed for: D) Donor chimerism presented at %CD45.2^+^YFP^+^ of each indicated HSPC population; E) Absolute numbers of donor-derived (CD45.2^+^YFP^+^) HSPC in the bone marrow from 1 femur and 1 tibia per mouse; F) Donor chimerism presented at %CD45.2^+^YFP^+^ of each indicated MP population; G) Granulocyte derived from donors in peripheral blood and bone marrow. Data are representative of 3 separate experiments with n=10 per group, presented as mean ± s.d. C) Unpaired Mann-Whitney u-test. D-G) Two-way ANOVA with Tukey’s multiple comparisons test. *, p<0.05; **, p<0.01; ***, p<0.001; ****, p<0.0001.

We subsequently treated the recipient mice ± IL-1β for 20 days (Fig. 2A). To confirm that our transplant conditions did not alter hematopoietic responses to IL-1β, we first assessed the hematopoietic output of wild-type (WT) competitor and recipient populations. As previously reported, chronic IL-1β increased production of myeloid-biased MPP3, GMP, and granulocytes (Fig. S1G-H, (Pietras et al., 2016)). We next evaluated competition dynamics within HSC, MPP, and MP by assessing donor chimerism within each compartment. Importantly, in the absence of chronic IL-1β, *Cebpa*^+/+^ and *Cebpa*^Δ/Δ^ chimerism was equivalent, suggesting the *Cebpa*^Δ/Δ^ LSK population does not possess a hematopoietic cell intrinsic competitive advantage. In contrast, chronic IL-1β administration triggered selective expansion of *Cebpa*^Δ/Δ^ ST-HSC, MPP2, MPP3 and MPP4 compartments (Fig. 2D). This expansion was also reflected in the absolute number of donor-derived *Cebpa*^Δ/Δ^ cells, with the largest numerical expansion within the MPP3 fraction (Fig. 2E). Chronic IL-1β-dependent expansion of donor-derived *Cebpa*^Δ/Δ^ cells likewise occurred in downstream Pre-MegE and Pre-GM populations (Fig. 2F). At this time point, we observed significant deficits in *Cebpa*^Δ/Δ^ donor-derived granulocytes in the periphery (Fig. 2G), consistent with a block in myeloid differentiation. By using less inflammatory methods for transplant, we did not see a hematopoietic cell intrinsic competitive advantage of *Cebpa*^Δ/Δ^ LSK cells. On the other hand, our data provide evidence that chronic inflammatory signaling via IL-1β triggers selective expansion of *Cebpa*^Δ/Δ^ HSPC.

### Chronic IL-1β-driven myeloid gene programs are *Cebpa*-dependent

We found that the *Cebpa*^Δ/Δ^ MPP3 compartment exhibited the most striking expansion during chronic IL-1β exposure. Because chronic IL-1β and *Cebpa* deficiency exert opposing effects on myeloid lineage output, we chose to investigate the molecular mechanism by which *Cebpa*^Δ/Δ^ confers a fitness advantage upon the GM-biased MPP3 in the context of chronic IL-1β exposure. Thus, we isolated both donor and competitor-derived MPP3 from recipient mice and performed RNA sequencing (RNA-seq) (Fig. 3A).

**Figure 3:**
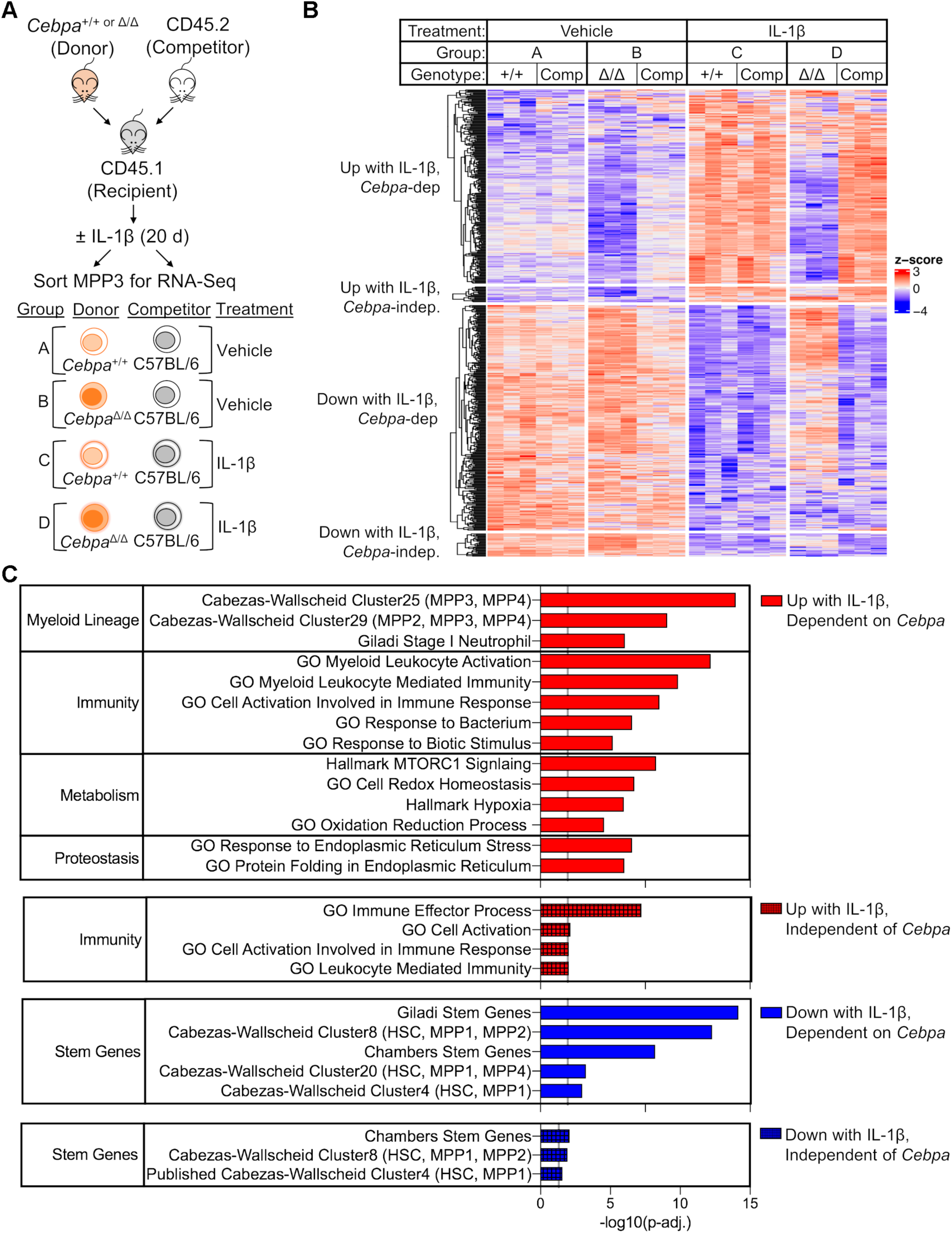
*Cebpa* mediates many of the chronic IL-1β gene expression changes. A) Experimental design for RNA-Seq (n=3 of 3-4 mice pooled in order to obtain 2000 donor (CD45.2^+^YFP^+^) or competitor (CD45.2^+^YFP^−^) MPP3). B) Heatmap of all genes significantly changed (p-adj. <0.05) by chronic IL-1β, grouped based on pattern of expression: Up with IL-1β, *Cebpa*-dependent; Up with IL-1β, *Cebpa*-independent; Down with IL-1β, *Cebpa*-dependent; Down with IL-1β, *Cebpa*-independent (genes for heatmap are provided in Table S1). C) Table showing select significant pathways (p-adj.<0.05) from over-representation analysis (ORA).

After validating *Cebpa* knockout by assessing its transcript levels in donor-derived MPP3 (Fig. S2A), we first determined the gene expression consequences of *Cebpa* deficiency in MPP3 (Fig. S2B). There were 494 differentially expressed genes (adjusted p-value, p-adj. <0.05) between *Cebpa*^Δ/Δ^ vs. *Cebpa*^+/+^ MPP3 isolated from vehicle-treated mice (Fig. S2B; gene list in Table S1). Chronic IL-1β had little effect on expression of these genes suggesting that *Cebpa* deficiency is epistatic to chronic IL-1β. We then focused on chronic IL-1β-driven gene expression changes by comparing gene expression in *Cebpa*^+/+^ MPP3 isolated from recipient mice treated ± chronic IL-1β. There were 555 differentially expressed genes (p-adj. <0.05). Interestingly, chronic IL-1β failed to induce most gene expression changes in *Cebpa*-deficient MPP3, suggesting that *Cebpa* is required for the majority of chronic IL-1β-induced gene programs in MPP3 (Fig. 3B). Only a small subset of chronic IL-1β-induced gene expression changes were independent of *Cebpa* (Fig. 3B). *Il1r1* expression was unchanged in *Cebpa*^Δ/Δ^ MPP3, ruling out the lack of IL-1 signaling as an explanation for the attenuated transcriptional signature (Fig. S2A). Consistent with this, *Nfkbia*, a canonical IL-1 target gene (Weber et al., 2010), is significantly downregulated in MPP3 isolated from chronic IL-1β-treated mice regardless of genotype (Fig. S2A).

To gain insight into the biological impact of these gene expression changes we performed over-representation analysis (ORA) (Fig. 3C). From the chronic IL-1β up-regulated genes that are dependent on *Cebpa*, 61 pathways were over-represented with p-adj. <0.05 (Fig. 3C), with many related to myeloid lineage or immunity. In accordance with ORA, pairwise gene set enrichment analyses (GSEA) revealed that *Cebpa* deficiency prevented activation of myeloid gene programs despite chronic IL-1β exposure (Fig. 4A-C, Fig. S2C). These data are consistent with the failure of donor-derived *Cebpa*^Δ/Δ^ to overproduce granulocytes in response to chronic IL-1β (Fig. 2G).

**Figure 4:**
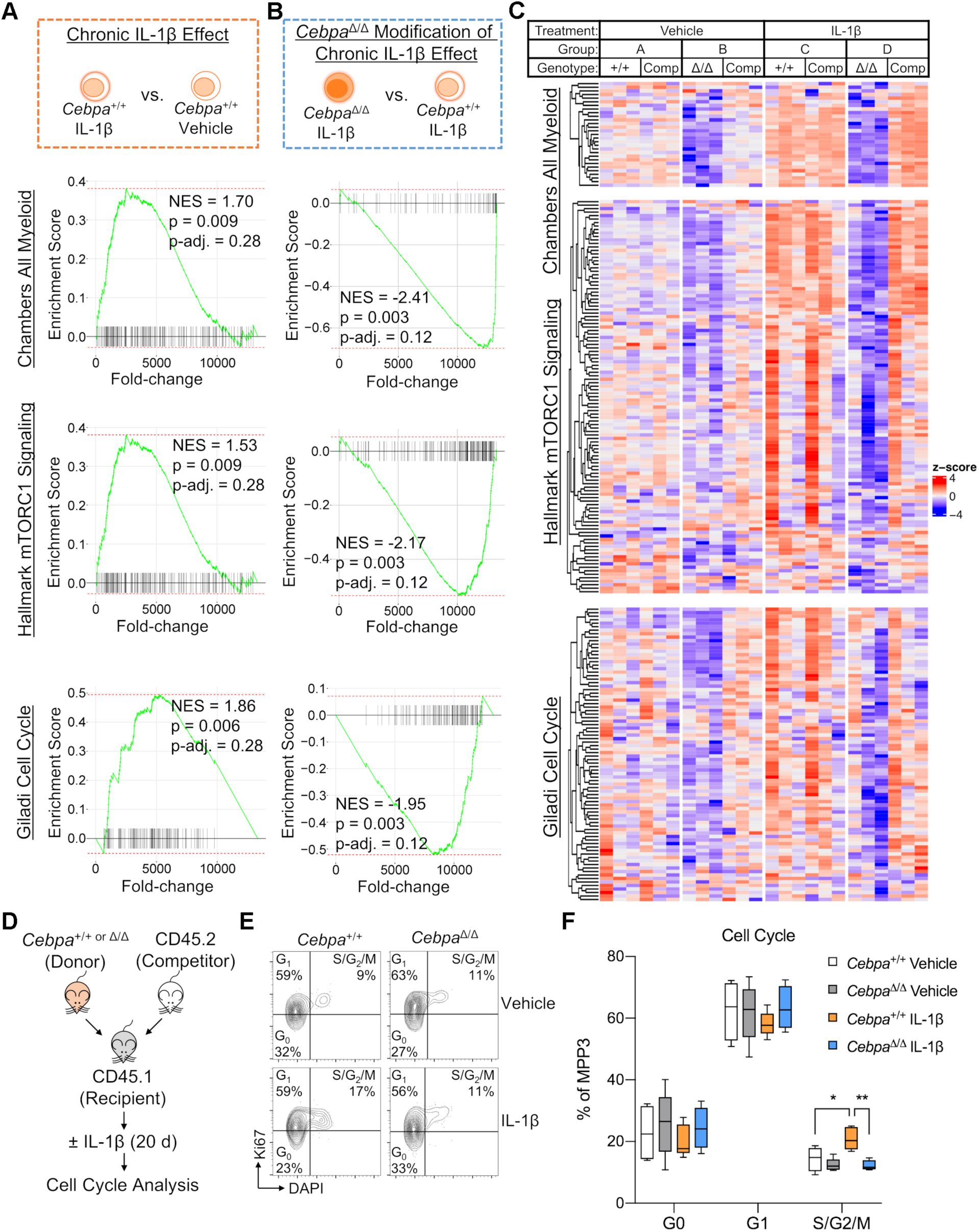
*Cebpa* deficiency counteracts chronic IL-1β-driven transcriptional programs. A-C) GSEA using “Chambers All Myeloid”, “Hallmark mTORC1 Signaling”, and “Giladi Cell Cycle” gene sets: A) Schematic and enrichment plots for pair-wise comparison between *Cebpa*^+/+^ MPP3 isolated from chronic IL-1β-treated vs. vehicle-treated mice (“Chronic IL-1β Effect”); B) Schematic and enrichment plots for pair-wise comparison between *Cebpa*^Δ/Δ^ vs. *Cebpa*^+/+^ MPP3 isolated from chronic IL-1β-treated mice (“*Cebpa*^Δ/Δ^ Modification of Chronic IL-1β Effect”); C) Heatmaps for leading edge genes from B (genes for heatmap are provided in Table S1). D) Experimental design for cell cycle analysis (n=5 per group). E) Representative flow cytometry plots for Ki67/DAPI cell cycle analysis. F) Graph of frequency of MPP3 in G0, G1, S/G2/M. D-F) Cell cycle data presented as mean ± s.d. Two-way ANOVA with Tukey’s multiple comparisons test. *, p<0.05; **, p<0.01.

Additional pathways that chronic IL-1β up-regulates in a *Cebpa*-dependent manner included those related to metabolism, protein synthesis, and redox homeostasis (Fig. 3C, Fig. 4A-C, Fig. S2C). Multiple studies have shown that tight regulation of these pathways is required for HSC function (Garcia-Prat et al., 2017; Ito and Suda, 2014; Signer et al., 2014). For instance, mTORC1 signaling, which is an important regulator of cell growth by regulating protein synthesis and cellular metabolism, has been shown to be important for myeloid cell differentiation (Peng et al., 2018; Zhang et al., 2019) and its hyperactivation can lead to HSC exhaustion (Gan and DePinho, 2009). HSPC differentiation is associated with increased metabolic and biosynthetic demands, so it seems appropriate that chronic IL-1β priming of MPP3 for myeloid differentiation is associated with up-regulation of the cellular processes required for differentiation. It was surprising that these pathways were also *Cebpa*-dependent, but this underscores the role of *Cebpa* as a master regulator coordinating not only myeloid gene programs, but also gene programs for cellular processes required to support differentiation.

### Chronic IL-1β-enhanced *Cebpa*-deficient MPP3 expansion is not due to changes in cell cycle activity

IL-1β can stimulate HSPC proliferation (Pietras et al., 2016). Conversely, C/EBPα inhibits cell cycle entry through direct interaction with CDKs and E2F (Johnson, 2005), and *Cebpa*-deficient HSC exhibit loss of quiescence (Hasemann et al., 2014; Ye et al., 2013). Thus, we reasoned that chronic IL-1β may directly increase *Cebpa*^Δ/Δ^ MPP3 proliferation, resulting in their selective expansion. Unexpectedly, *Cebpa* deficiency prevented chronic IL-1β-driven upregulation of cell cycle genes (Fig. 4A-C). Likewise, cell cycle analysis of donor-derived *Cebpa*^+/+^ and *Cebpa*^Δ/Δ^ MPP3 isolated from recipient mice treated ± chronic IL-1β revealed a modest but significant increase in the frequency of *Cebpa*^+/+^ MPP3 in S/G2/M phase following treatment with chronic IL-1β, whereas cell cycle status of *Cebpa*^Δ/Δ^ MPP3 was unchanged (Fig. 4D-F), consistent with our gene expression data. Together, these data suggest that the competitive advantage of *Cebpa*^Δ/Δ^ MPP3 in the presence of IL-1β is not explained by changes in cell cycle activity.

### Chronic IL-1β-exposed *Cebpa*-deficient MPP3 activate a self-renewal gene program

To further assess the molecular basis for selective expansion of *Cebpa*^Δ/Δ^ MPP3 *in vivo* we analyzed genes downregulated by IL-1β in *Cebpa*^+/+^ but not *Cebpa*^Δ/Δ^ MPP3. ORA revealed that pathways over-represented with this expression pattern included those related to stemness (Fig. 3C), suggesting that chronic IL-1β represses stem cell gene programs in MPP3 via C/EBPα. GSEA supported these findings, showing enrichment for stem cell gene signatures in *Cebpa*^Δ/Δ^ MPP3 from chronic IL-1β-treated recipient mice (Fig. 5A-C, Fig. S2C). To validate these findings, we isolated donor-derived *Cebpa*^+/+^ and *Cebpa*^Δ/Δ^ MPP3 from recipient mice treated ± chronic IL-1β for a Fluidigm gene expression array and found that expression of genes essential for HSC self-renewal, such as *Bmi1, Foxo3, Mpl*, and *Mycn* (Iwama et al., 2004; Laurenti et al., 2008; Miyamoto et al., 2007; Nitta et al., 2020; Yoshihara et al., 2007), were not only preserved but elevated in *Cebpa*^Δ/Δ^ MPP3 as compared to *Cebpa*^+/+^ MPP3 from chronic IL-1β-exposed mice (Fig. 5D-E). Thus, the fitness advantage in the context of chronic IL-1β may be attributed to the coordinate increase in self-renewal-associated genes in *Cebpa*^Δ/Δ^ MPP3.

**Figure 5:**
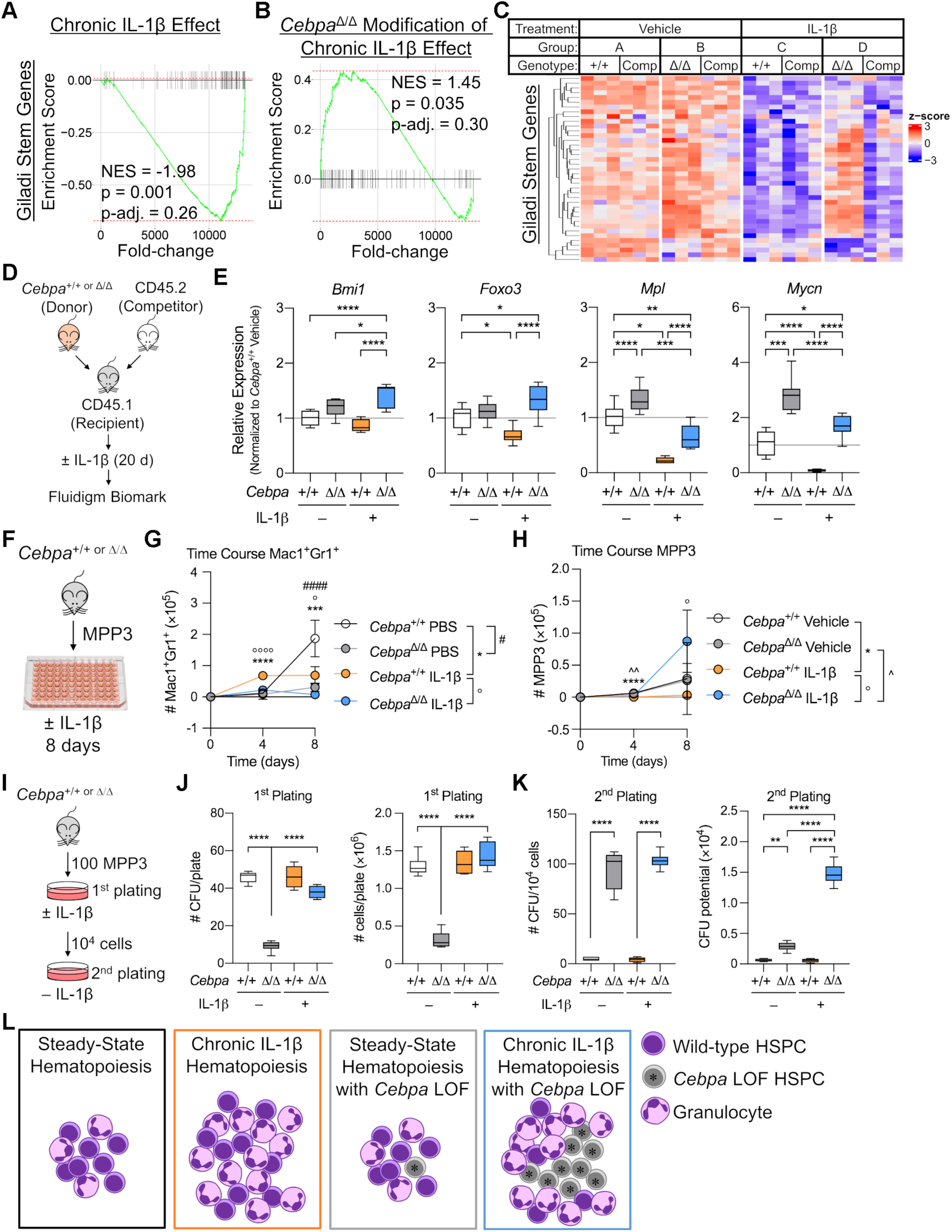
Chronic IL-1β triggers aberrant expansion potential of *Cebpa*-deficient MPP3. A-C) GSEA using “Giladi Stem Genes” gene set: A) Schematic and enrichment plot for pair-wise comparison between *Cebpa*^+/+^ MPP3 isolated from chronic IL-1β-treated vs. vehicle-treated mice (“Chronic IL-1β Effect”); B) Schematic and enrichment plot for pair-wise comparisons between *Cebpa*^Δ/Δ^ vs. *Cebpa*^+/+^ MPP3 isolated from chronic IL-1β-treated mice (“*Cebpa*^Δ/Δ^ Modification of Chronic IL-1β Effect”); C) Heatmap for leading edge genes from B (genes for heatmap are provided in Table S1). D) Experimental design for Fluidigm Biomark gene expression analysis (8 replicates from n=5 pooled mice per group). E) Relative expression of self-renewal genes (*Bmi1, Foxo3, Mpl*, and *Mycn*) calculated by first normalizing to *Gusb* within each group, then normalizing to *Cebpa*^+/+^ vehicle group. F-H) Liquid culture data representative of 3 separate experiments with n=3-5 per group: F) Experimental design for MPP3 liquid culture; G) Time course of Mac1^+^Gr1^+^ absolute numbers; H) Time course of MPP3 absolute numbers. I-K) CFU assay representative of 2 experiments with n=3 or 6 per group: I) Experimental design for MPP3 CFU; J) CFU counts per plate and total cell counts per plate from initial plating; K) CFU counts per plate upon re-plating 10^4^ cells and calculated cumulative CFU potential from 2^nd^ plating. L) Model of chronic-IL-1β-dependent *Cebpa*-deficient HSPC aberrant expansion potential. Data are presented as mean ± s.d. Two-way ANOVA with Tukey’s multiple comparisons test. *, p<0.05; **, p<0.01; ***, p<0.001; ****, p<0.0001, except in G and H where symbols for different comparisons are indicated.

### *Cebpa* deficiency selectively exacerbates chronic IL-1β gene expression programs in its competitors

Somatic cell fitness describes how cell intrinsic properties (*Cebpa* deficiency) interact with microenvironmental factors (IL-1β) and other cells within its niche. Cell competition dynamics are appreciated to be an important factor in tumorigenesis (Bowling et al., 2019). We hypothesized that WT competitor MPP3 would exhibit differential gene expression in competition against a more fit population (vs. *Cebpa*^Δ/Δ^ MPP3 in chronic IL-1β) versus in neutral competition (vs. *Cebpa*^+/+^ MPP3 in chronic IL-1β), even though they are identical by genotype and treatment. We performed GSEA pair-wise comparisons between WT competitor MPP3 transplanted with *Cebpa*^+/+^ or *Cebpa*^Δ/Δ^ in the context of chronic IL-1β (Fig. S2D). Interestingly, WT MPP3 that were competed against *Cebpa*^Δ/Δ^ MPP3 had enrichment for myeloid gene programs above that of WT MPP3 that were competed against *Cebpa*^+/+^ MPP3 in the context of chronic IL-1β. Further, WT MPP3 that were competed against *Cebpa*^Δ/Δ^ MPP3 were less enriched for stem gene programs as compared to WT MPP3 competed against *Cebpa*^+/+^ MPP3 in the context of chronic IL-1β (Fig. S2D). When the same comparison was made between competitors isolated from vehicle-treated mice, no comparisons were significantly different (Fig. S2E). This suggests that WT competitors differ by gene expression only in the presence of a fitness differential. These data provide evidence at the molecular level that competition in the presence of *Cebpa*^Δ/Δ^ MPP3 exacerbated chronic IL-1β activation of myeloid gene programs and repression of stem gene programs.

### Chronic IL-1β triggers aberrant expansion potential of *Cebpa*-deficient MPP3

To assess the direct impact of IL-1β on *Cebpa*^+/+^ or *Cebpa*^Δ/Δ^ MPP3 expansion, we sorted MPP3 from primary *Cebpa*^+/+^ or *Cebpa*^Δ/Δ^ mice for liquid culture ± IL-1β and assessed MPP3 maintenance and differentiation by flow cytometry (Fig. 5F, Fig. S2F). Similar to studies performed with purified HSC (Pietras et al., 2016), IL-1β induced precocious differentiation of *Cebpa*^+/+^ MPP3 and early accumulation of Mac1^+^Gr1^+^ cells. Despite IL-1β treatment, *Cebpa*^Δ/Δ^ MPP3 formed very few differentiated Mac1^+^Gr1^+^ cells over the 8-day culture (Fig. 5G). Consistent with our *in vivo* data, there was no difference in the number of phenotypic *Cebpa*^+/+^ or *Cebpa*^Δ/Δ^ MPP3 cells after 8 days of culture without IL-1β. However, IL-1β induced selective expansion of *Cebpa*^Δ/Δ^ MPP3 in culture (Fig. 5H). We also analyzed serial CFU activity of *Cebpa*^+/+^ or *Cebpa*^Δ/Δ^ MPP3 ± IL-1β (Fig. 5I). As seen with LSK cells, IL-1β did not affect the total number of CFUs or cells formed by *Cebpa*^+/+^ MPP3 on the initial round of plating. While *Cebpa*^Δ/Δ^ MPP3 formed fewer CFU and had fewer total cells in the absence of IL-1β, their clonogenic activity increased significantly when cultured with IL-1β (Fig. 5J). Upon replating, *Cebpa*^+/+^ cells formed very few CFU, but *Cebpa*^Δ/Δ^ cells formed significantly more CFU regardless of whether previously plated with or without IL-1β (Fig. 5K), and *Cebpa*^Δ/Δ^ MPP3 retained clonogenic potential up to the 3^rd^ and 4^th^ replatings (Fig. S2G). Again, we calculated the cumulative CFU potential at replating and found that IL-1β significantly increased the cumulative CFU potential of *Cebpa*^Δ/Δ^ MPP3 (Fig. 5E). This demonstration of IL-1β-specific *Cebpa*^Δ/Δ^ MPP3 expansion in cell culture recapitulates our *in vivo* data and further isolates the fitness advantage as reliant specifically on IL-1β. In addition, these data suggest that *Cebpa*^Δ/Δ^ MPP3 expand following IL-1β exposure via triggering aberrant self-renewal activity, which is consistent with upregulation of key stem gene *in vivo*.

In conclusion, using a mouse model with HSPC-specific *Cebpa* deletion and less inflammatory competitive transplant methods, we show that *Cebpa*-deficient HSPC do not have a hematopoietic cell intrinsic competitive advantage relative to WT HSPC. Strikingly, chronic IL-1β exposure triggered a fitness advantage for *Cebpa*-deficient MPPs. While chronic IL-1β remodels the HSPC compartment by expanding myeloid-biased MPP to increase myeloid output, C/EBPα blocks myeloid differentiation. Together, these phenotypes cooperate and ultimately result in chronic IL-1β-dependent expansion of *Cebpa*^Δ/Δ^ cells. While prior studies had shown an inherent competitive advantage of *Cebpa*^Δ/Δ^ HSC, these investigations used polyinosinic:polycytidylic acid to induce Cre recombination and lethal irradiation to condition transplant recipients (Ye et al., 2013; Zhang et al., 2004). Both approaches are highly inflammatory and impair BM function. In addition, *Mx1-*Cre-mediated *Cebpa* deletion rapidly depleted granulocytes, which can activate MPP proliferation (Sawen et al., 2016) and confound functional studies of HSPC. Our data demonstrate that an inflammatory environment is required to reveal the competitive advantage of *Cebpa*-deficient HSPC.

Transcriptional analyses revealed that *Cebpa* deficiency prevents chronic IL-1β-driven activation of myeloid differentiation, as well as metabolic and proteostatic gene programs important for differentiation. Despite the pre-existing molecular priming of MPP3 toward myeloid lineage differentiation (Cabezas-Wallscheid et al., 2014; Pietras et al., 2015), our data show chronic IL-1β further activates these pathways, consistent with the capacity of myeloid lineage progenitors to overproduce myeloid cells in response to ‘emergency’ cues (Herault et al., 2017; Kang et al., 2020; Manz and Boettcher, 2014). The dependency of these gene programs on C/EBPα is consistent with its role as a key master regulator that initiates myeloid lineage priming in the LSK population (Pundhir et al., 2018). It is interesting that chronic IL-1β has been shown to induce precocious myeloid differentiation via PU.1 (Pietras et al., 2016). C/EBPα can regulate PU.1 expression and enhancer binding in pre-GMs (Pundhir et al., 2018). In fact, PU.1 expression was significantly reduced in *Cebpa*^Δ/Δ^ MPP3 in the context of chronic IL-1β (Fig S2A). Collectively, these data suggest that *Cebpa* is important for GM-lineage specification in MPP3, and that chronic IL-1β primes GM-lineage output via *Cebpa*-dependent gene programs in MPP3. In addition, *Cebpa* deficiency prevents chronic IL-1β-driven repression of stem cell self-renewal genes, *Bmi1, Foxo3, Mpl*, and *Mycn* (Iwama et al., 2004; Laurenti et al., 2008; Miyamoto et al., 2007; Nitta et al., 2020; Yoshihara et al., 2007), and C/EBPα has been demonstrated to directly bind to the promoters of *Mycn* (Ye et al., 2013) and *Foxo3* (Hasemann et al., 2014). This is associated with expansion of immature HSPC with aberrant serial replating potential from *Cebpa*-deficient MPP3 following IL-1β treatment.

While steady-state hematopoiesis results in balanced lineage output, chronic IL-1β expands the MPP compartment to increase myeloid output. However, IL-1β-induced differentiation gene programs likely serve as a limiting mechanism that prevents aberrant accumulation of HSPC. Our data support a model whereby aberrant expansion of HSPC with *Cebpa* loss-of-function (LOF) is an emergent property triggered by inflammatory signals such as chronic IL-1β. Our findings suggest *Cebpa* LOF HSPC fail to differentiate and instead activate expression of stem cell self-renewal genes, thereby promoting their selective expansion while WT HSPC differentiate in response to chronic IL-1β (Fig. 5L).

There is growing evidence of context-specific expansion of oncogenically-initiated hematopoietic progenitors: *BCR-ABL* or *Ppm1d*-deficiency with chemotherapy (Bilousova et al., 2005; Hsu et al., 2018; Kahn et al., 2018); *Notch* activation, *Tp53*-deficiency, or *Cebpa*-deficiency following γ-irradiation (Bondar and Medzhitov, 2010; Fleenor et al., 2015b; Marusyk et al., 2009; Marusyk et al., 2010); BCR-ABL, *Myc, NRas*^*V12*^, or AML1-ETO in aging (Henry et al., 2015; Henry et al., 2010; Vas et al., 2012); and *Jak*^*V617F*^ with TNF-α exposure (Fleischman et al., 2011) or *Tet2*-knockout with LPS challenge (Cai et al., 2018). Here, we show that chronic IL-1β is sufficient to drive selective expansion of *Cebpa*-deficient MPPs, directly connecting an inflammatory signaling pathway to a transcriptional fate network. These findings have important implications for the earliest events that could drive CHIP, MDS, MPN or AML. Initially, IL-1 drives selection for mutations in HSPC that provide resistance to the pro-differentiative effects of IL-1. This expansion will proportionally increase the odds of successive mutation accumulation in the same clone, increasing the possibility of leukemic transformation. Once transformed, IL-1 can then act as a growth factor for AML blasts (Carey et al., 2017) and contribute to more aggressive disease (Barreyro et al., 2012). Therefore, inflammation could be an important driver of leukemogenesis and could represent a potential target for intervention.

## Materials and Methods

### Mice

*Cebpa*^loxP/loxP^ mice were previously described (Zhang et al., 2004). *HSC-SCL-Cre-ER*^*T*^ transgenic mice were generously provided by Joachim Gothert (Gothert et al., 2005). *R26R-EYFP* transgenic reporter mice were generously provided by Dennis Roop (Srinivas et al., 2001). All mice were bred to be heterozygous for *R26R-EYFP* and heterozygous for *HSC-SCL-Cre-ER*^*T*^, while each litter contained a mixture of *Cebpa* genotypes. To induce recombination, mice were treated intraperitoneally (i.p.) with 2.5 mg of tamoxifen (prepared at 10 mg/mL in corn oil) daily for 3 days and were used for experiments 7 days after the final dose of tamoxifen. Genotyping of mice and detection of recombined product were genotyped as previously described (Zhang et al., 2004). C57BL/6 (Ly5.2) and congenic (Ly5.1) mice were ordered from Charles Rivers Laboratories. All procedures were performed in accordance with the University of Colorado Anschutz Medical Campus Institutional Animal Care and Use Committee (IACUC)-approved animal protocols.

### Cell Preparation and Flow Cytometry

Peripheral blood for flow cytometry analysis or complete blood count was collected in heparin (50 units/mL) from the sub-mandibular vein from live mice or via cardiac puncture from terminal mice. CBC analyses were run on a Heska HT5 blood analyzer. For flow cytometry analyses, red blood cells (RBCs) were hemolyzed with Ammonium-Chloride-Potassium (ACK) lysis buffer, and cells were washed and filtered. Cell were stained for mature immune cells using the following antibodies: rat IgG for blocking, Mac1 (M1/70) Brilliant Ultraviolet™ (BUV) 395, Gr1 (Rb6-8C5) Pacific Blue™ (PB), CD45.2 (104) biotin, streptavidin Brilliant Violet™ (BV)605, B220 (RA3-6B2) BV786, CD45.1 (A20) PerCP-Cy-5.5, CD8 (53-6.7) PE, Ter-119 (TER-119) PE-Cy5, CD4 (GK1.5) PE-Cy7, IgM (RMM-1) APC, CD3 (17A2) Alexa Fluor® (AF) 700, and CD19 (1D3) APC-Cy7. Bone marrow for flow analysis was obtained by flushing cells from one tibia and one femur per mouse. RBCs were hemolyzed with ACK lysis buffer, and cells were washed and filtered. Cells counts were collected on the Guava® easyCyte™ System (Millipore). Bone marrow cells were stained for mature immune cells as with peripheral blood and for hematopoietic stem and progenitor cells (HSPC) with the following antibodies: rat IgG, Mac1 BUV395, Sca1 (D7) BV421, CD41 (MWReg30) BV510, CD45.2 biotin, streptavidin BV605, CD105 (MJ7/18) BV711, CD150 (TC15-12F12.2) BV785, CD45.1 PerCP-Cy.5.5, Flk2 (A2F10) PE, lineage cocktail (B220, CD3, CD4, CD5 (53-7.3), CD8, Gr1, Ter119) conjugated with PE-Cy5, FcgR (93) PE-Cy7, CD34 (RAM34) eFluor™(e)660, CD48 (HM48-1) AF700, cKit (2B8) APC-Cy7. Data were collected on the ZE5 cell analyzer (Biorad) or the LSRFortessa™ (BD Biosciences) and analyzed with FlowJo™ Software v10. Bone marrow for cell sorting was obtained by crushing all long bones and spines from each mouse. RBCs were hemolyzed with ACK lysis buffer, and cells were washed and filtered. Bone marrow cells were lineage depleted using Mouse Lineage Cell Depletion Kit (Miltenyi Biotec). Cells were stained to identify HSPC with the following antibodies: rat IgG, CD48 e450, CD45.2 biotin, streptavidin BV605, CD150 BV786, CD45.1 PerCP-Cy5.5, Flk2 PE, cKit PE-Cy7, Sca1 APC, and lineage cocktail conjugated with APC-Cy7. Cell sorting was performed with the 100 µm nozzle on the FACSAria™ Fusion (BD Biosciences). Flow cytometry analysis was performed on cells from liquid culture and on individual colonies from methylcellulose with the following antibodies: rat IgG, Mac1 BUV395, Gr1 PB, Sca1 APC, and cKit PE-Cy7. Data were collected on the ZE5 cell analyzer (Biorad) and analyzed with FlowJo™ Software v10.

### Cell Culture

For liquid cultures, 200 YFP^+^ LSK cells were sorted from *Cebpa*^+/+^ or *Cebpa*^Δ/Δ^ mice into wells of a 96-well culture plate. Cells were grown in IMDM (Gibco) with 20% heat-inactivated FBS (HyClone), 50 μM β-mercaptoethanol, IL-6 (100 ng/mL), Flt3-L (100 ng/mL), SCF (100 ng/mL), and IL-11 (10 ng/mL) as previously described (Varnum-Finney et al., 2003). For colony-forming unit (CFU) assays, 100 YFP^+^ LSK cells were sorted from *Cebpa*^+/+^ or *Cebpa*^Δ/Δ^ mice and plated in methylcellulose (R&D HSC006; 1 mL in 3 cm dish) containing IMDM, antibiotic-antimycotic (Gibco), SCF (25 ng/mL), Flt3-L (25 ng/mL), IL-11 (25 ng/mL), IL-3 (10 ng/mL), GM-CSF (10 ng/mL), TPO (25 ng/mL), EPO (4 U/mL) with or without IL-1β (25 ng/mL). Colonies were counted and scored under a dissecting scope after 7 days. For serial re-plating, 10^4^ cells were re-plated in fresh methylcellulose with or without IL-1β. All cytokines were obtained from Peprotech. We calculated cumulative CFU potential with the equation below:

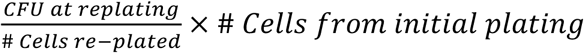

### Competitive Transplants

To obtain a starting donor chimerism of ∼5%, 2×10^3^ LSK cells were sorted and transplanted with 2×10^6^ competitor whole BM cells into mildly conditioned congenic mice. Mild conditioning was achieved by treating mice with 20 mg/kg of busulfan 4 days before transplant and 30 µg of anti-CD4 antibody, clone GK1.5 (BioXcell) and 30 µg of anti-CD8 antibody, clone 2.43 (Bio-X-Cell) 2 days before transplant. Busulfan was prepared fresh at 20 mg/mL in DMSO and diluted in saline to 2 mg/mL, while kept warm and protected from light. CD4 and CD8 T cell depleting antibodies were prepared at 30 µg per 0.1 mL of PBS. Initial chimerism was assessed 3 weeks post-transplant, and mice were subsequently treated daily for 20 days with 0.1% BSA/PBS or 0.5 µg IL-1β (Peprotech) in 0.1% BSA/PBS via i.p. injection.

### Fluidigm Biomark Gene Expression Analysis

Cells were sorted into 96-well plates at 50 cells/well in 5 µL CellsDirect™ 2× Reaction Buffer (Invitrogen). Cells lysates were reverse transcribed using Superscript™ III (Invitrogen) and pre-amplified for 20 rounds using a custom-designed set of target-specific primers. Pre-amplified cDNA was treated with Exonuclease I (New England Biolabs) to remove excess primers, diluted in DNA suspension buffer (Clontech), and loaded with custom-designed primer sets onto a Fluidigm 96.96 DynamicArray IFC and run on a BioMark System (Fluidigm) using SsoFast Sybr Green for detection (Bio-Rad). Data were analyzed by the ΔΔCt method using Fluidigm software and normalized to *Gusb* expression.

### RNA Sequencing

Cells were sorted directly into RLT Buffer and RNA was extracted using RNeasy Micro Kit (Qiagen). Library construction was performed using the SMARTer® Stranded Total RNA-Seq Kit v2 – Pico Input Mammalian (Takara), and paired-end sequencing was performed on the NovaSeq 6000 instrument (Illumina) by the University of Colorado Cancer Center Genomics and Microarray Core. Illumina adapters were removed using BBDuk (Bushnell, 2019) and reads <50bp after trimming were discarded. Reads were aligned and quantified using STAR (2.6.0a) (Dobin et al., 2012) to the Ensembl mouse transcriptome (GRCm38.p6 release 96). Normalization and differential expression were calculated using the limma R package (Ritchie et al., 2015). Differentially expressed genes (adjusted p. value < 0.05) comparing the donors of group C versus the donors of group A were determined to be influenced by IL-1β (Fig. 3A). These genes were further determined to be *Cebpa*-dependent or -independent based on whether they were significantly altered when comparing the donors of group D to the donors of group B. Over-representation analysis was performed on these genes using the ClusterProfiler R package (Yu et al., 2012) with Hallmark and GO Biological Processes gene sets from the Molecular Signatures Database (Liberzon et al., 2011) along with select published gene sets (Cabezas-Wallscheid et al., 2014; Chambers et al., 2007; Giladi et al., 2018; Ye et al., 2013). Gene set enrichment analysis (GSEA) was performed on the fold change of the genes for each comparison using the fGSEA R package (Sergushichev, 2016) with the same gene sets. Heatmaps were generated with the ComplexHeatmap R package (Gu et al., 2016) following z-score transformation. The genes in the pathway heatmaps were selected because of their presence in the leading edge as determined by fGSEA for the indicated comparison.

### Cell Cycle Analysis

Cells were sorted and fixed into FACS buffer, then immediately fixed in BD Cytofix/Cytoperm. Cells were re-stained with surface stain, and intracellular staining for Ki67 (SolA15) APC was performed according to the Fixation/Permeabilization Solution Kit (BD Bioscience) and as previously described (Jalbert and Pietras, 2018). Data were collected on the ZE5 cell analyzer (Biorad) and analyzed with FlowJo™ Software v10.

### Statistics

Unpaired Mann-Whitney u-test (for *Cebpa*^+/+^ vs *Cebpa*^Δ/Δ^ comparison) or two-way ANOVA with Tukey’s multiple comparisons test (for combined *Cebpa*^+/+^ vs *Cebpa*^Δ/Δ^ vehicle vs. chronic IL-1β comparison) was performed in Prism Software v8 (GraphPad). *, p<0.05; **, p<0.01; ***, p<0.001; ****, p<0.0001.

## Online Supplemental Material

Fig. S1 shows that HSPC-specific knockout of *Cebpa* causes a defect in myelopoiesis and that chronic IL-1β-enhanced myeloid output is retained using our transplant conditions. Fig. S2 shows that *Cebpa* deficiency is epistatic to chronic IL-1β, by preventing upregulation of myeloid differentiation programs and downregulation of stem gene programs, ultimately resulting in IL-1β-selective expansion of *Cebpa*-deficient MPP3. Table S1 compiles lists of genes from heatmaps presented in Fig. 3B, Fig. 4C, Fig. 5C, and Fig. S2B.

## Acknowledgments

We thank Patricia Ernst, Craig Jordan, and Marco De Dominici for critical review of the manuscript, and Pavel Davizon-Castillo for performing CBCs. These studies were supported by grants from the National Institutes of Health R01-AG067548 and R01-DK119394 to J.D. and E.M.P., R01-CA180175 and U01-AG066099 to J.D., R35-CA197697 and P01-HL131477 to D.G.T., F30-CA210383 and T32-AG000279 to K.C.H., F31-HL138754 to J.L.R., the Cleo Meador and George R. Scott Endowed Chair of Medicine in Hematology to E.M.P., Leukemia and Lymphoma Society SCOR grant 7020-19 to J.D., and the Courtenay C. and Lucy Patten Davis Endowed Chair in Lung Cancer Research to J.D. We thank the University of Colorado Cancer Center Shared Resources (Flow Cytometry, Genomics and Bioinformatics/Biostatistics) supported by P30-CA046934, the Human Immune Monitoring Shared Resource, and the Clinimmune Flow Core.

The authors have no conflicting financial interests.

## Author Contributions

K.C.H performed all experiments with assistance from J.S.C., V.Z., and J.L.R. K.C.H and A.G. analyzed and interpreted RNA-seq data. K.C.H, E.M.P., and J.D. designed research studies, interpreted results, and wrote the manuscript.

---

**Figure S1:**
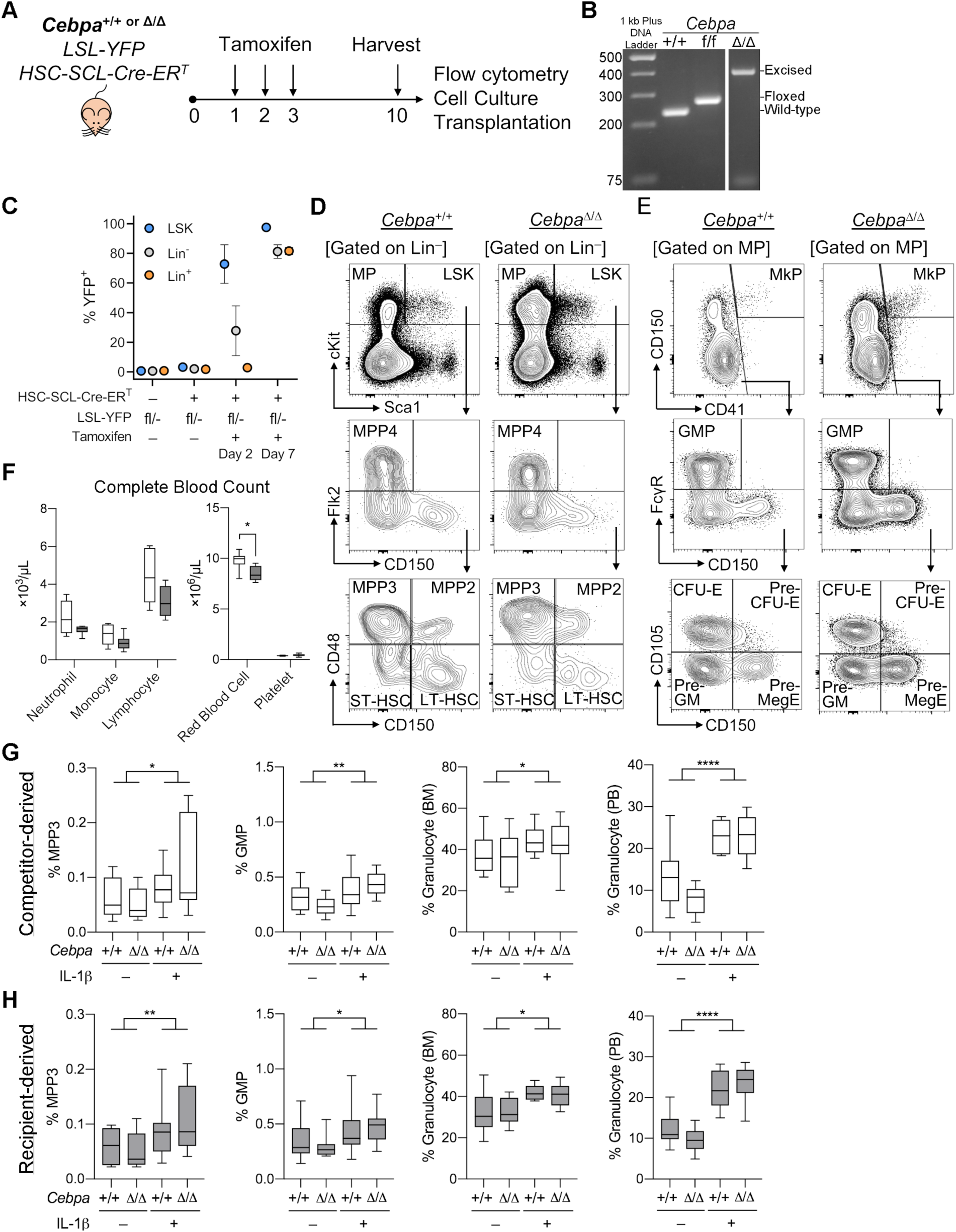
Myeloid differentiation in primary *Cebpa*-deficient mice and recipient/competitor HSPC. A) Schematic to achieve traceable, tamoxifen-inducible, HSPC-specific knockout of *Cebpa*. B) PCR products to detect wild-type (235 bp), floxed (269 bp), or excised allele (361 bp) run on a 2% agarose gel with 1 kb Plus DNA ladder (Thermo Fisher). C) Kinetics of tamoxifen-induced YFP expression within Lin^−^Sca1^+^cKit^+^ (LSK), Lin^−^, and Lin^+^ BM fractions. Tamoxifen did not influence HSPC behavior in cell culture or IL-1 levels in BM fluid (data not shown). D-F) Hematopoietic characterization of *Cebpa*^+/+^ and *Cebpa*^Δ/Δ^ mice (n=9 each) 7 days post-deletion: D) Representative flow cytometry plots to characterize HSPC populations; E) Representative flow plots to characterize MP; F) Complete blood count (CBC) of peripheral blood from *Cebpa*^+/+^ and *Cebpa*^Δ/Δ^ mice. G-H) Validation of chronic IL-1β phenotype in competitive transplants by characterizing hematopoietic output: G) MPP3, GMP, and granulocytes derived from competitors; H) MPP3, GMP, and granulocytes derived from recipients. Presented as mean ± s.d. F) Unpaired Mann-Whitney u-test used to analyze data. G-H) Two-way ANOVA with Tukey’s multiple comparisons test. *, p<0.05; **, p<0.01; ***, p<0.001; ****, p<0.0001.

**Figure S2:**
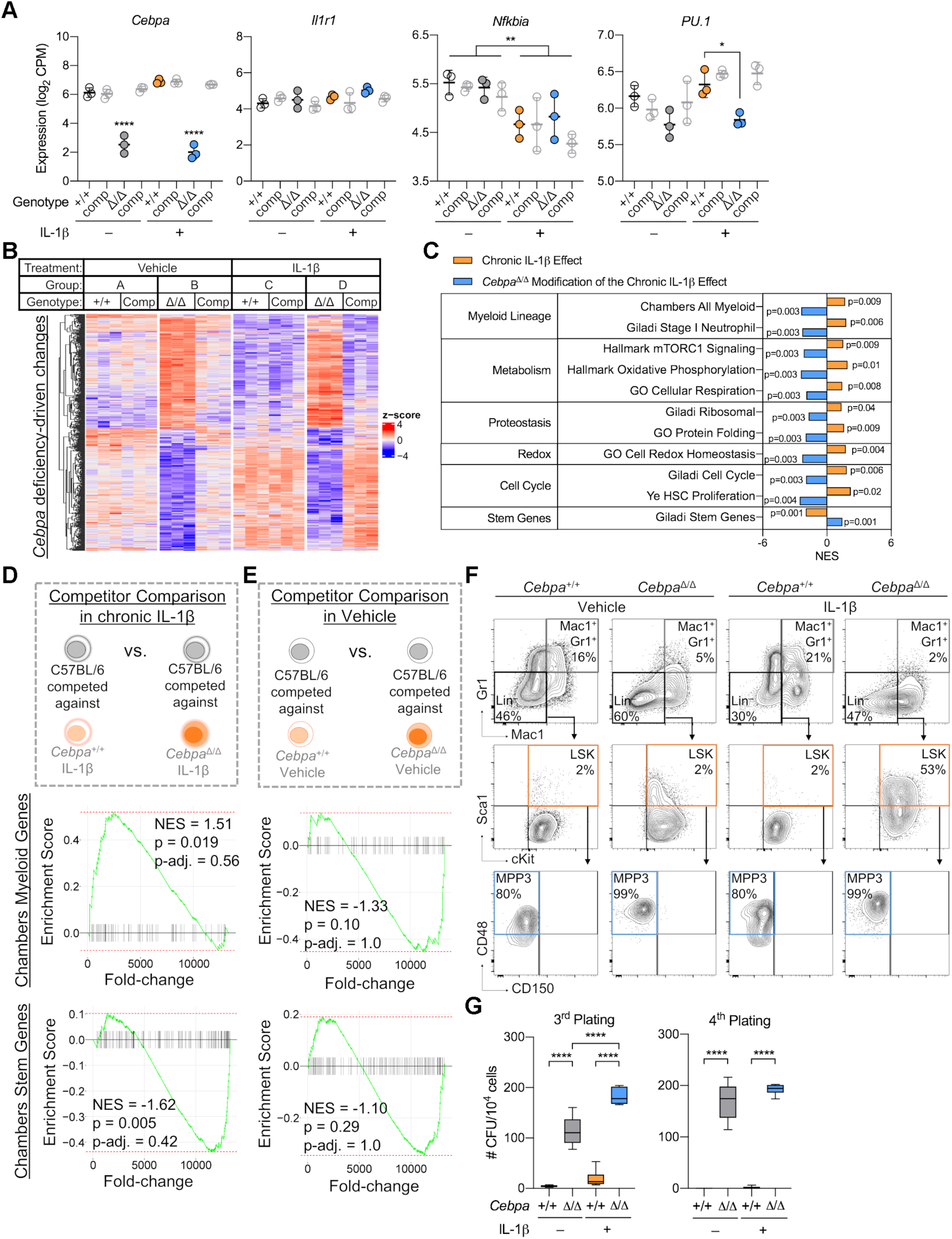
*Cebpa* deficiency confers a fitness advantage in the context of chronic IL-1β *in vivo*. A) Expression of *Cebpa, Il1r1, Nfkbia* and *Spi1* plotted as log_2_(CPM) from RNA-seq. B) Heatmap of all genes significantly changed (p-adj. <0.05) by *Cebpa* deficiency (genes for heatmap are provided in Table S1). C) Table summarizing GSEA Normalized Enrichment Scores (NES) for select gene sets for “Chronic IL-1β Effect” and “*Cebpa*^Δ/Δ^ Modification of Chronic IL-1β Effect” pair-wise comparisons. D-E) GSEA with “Chamber All Myeloid” and “Chambers Stem Genes” gene sets: D) Schematic and enrichment plots for pair-wise comparison between C57BL/6 MPP3 competed against *Cebpa*^Δ/Δ^ or *Cebpa*^+/+^ from chronic IL-1β-treated mice; E) Schematic and enrichment plots for pair-wise comparison between C57BL/6 MPP3 competed against *Cebpa*^Δ/Δ^ or *Cebpa*^+/+^ from vehicle-treated mice. F) Representative flow cytometry plots for liquid culture analyses. G) 3^rd^ and 4^th^ plating CFU counts per plate upon re-plating 10^4^ cells. Data are presented as mean ± s.d. Two-way ANOVA with Tukey’s multiple comparisons test. *, p<0.05; **, p<0.01; ***, p<0.001; ****, p<0.0001.

**Table S1:** Table of gene names from heat maps in Fig. 3B, Fig. 4C, Fig. 5C, and Fig. S2B.

## References

Barreyro, L., B. Will, B. Bartholdy, L. Zhou, T.I. Todorova, R.F. Stanley, S. Ben-Neriah, C. Montagna, S. Parekh, A. Pellagatti, J. Boultwood, E. Paietta, R.P. Ketterling, L. Cripe, H.F. Fernandez, P.L. Greenberg, M.S. Tallman, C. Steidl, C.S. Mitsiades, A. Verma, and U. Steidl. 2012. Overexpression of IL-1 receptor accessory protein in stem and progenitor cells and outcome correlation in AML and MDS. Blood 120:1290–1298.

Bereshchenko, O., E. Mancini, S. Moore, D. Bilbao, R. Mansson, S. Luc, A. Grover, S.E. Jacobsen, D. Bryder, and C. Nerlov. 2009. Hematopoietic stem cell expansion precedes the generation of committed myeloid leukemia-initiating cells in C/EBPalpha mutant AML. Cancer Cell 16:390–400.

Bilousova, G., A. Marusyk, C.C. Porter, R.D. Cardiff, and J. DeGregori. 2005. Impaired DNA replication within progenitor cell pools promotes leukemogenesis. PLoS Biol 3:e401.

Bondar, T., and R. Medzhitov. 2010. p53-mediated hematopoietic stem and progenitor cell competition. Cell Stem Cell 6:309–322.

Bowling, S., K. Lawlor, and T.A. Rodriguez. 2019. Cell competition: the winners and losers of fitness selection. Development 146:

Bowman, R.L., L. Busque, and R.L. Levine. 2018. Clonal Hematopoiesis and Evolution to Hematopoietic Malignancies. Cell Stem Cell 22:157–170.

Busch, K., K. Klapproth, M. Barile, M. Flossdorf, T. Holland-Letz, S.M. Schlenner, M. Reth, T. Hofer, and H.R. Rodewald. 2015. Fundamental properties of unperturbed haematopoiesis from stem cells in vivo. Nature 518:542–546.

Bushnell, B. 2019. BBMap. In.

Cabezas-Wallscheid, N., D. Klimmeck, J. Hansson, D.B. Lipka, A. Reyes, Q. Wang, D. Weichenhan, A. Lier, L. von Paleske, S. Renders, P. Wunsche, P. Zeisberger, D. Brocks, L. Gu, C. Herrmann, S. Haas, M.A.G. Essers, B. Brors, R. Eils, W. Huber, M.D. Milsom, C. Plass, J. Krijgsveld, and A. Trumpp. 2014. Identification of regulatory networks in HSCs and their immediate progeny via integrated proteome, transcriptome, and DNA methylome analysis. Cell Stem Cell 15:507–522.

Cai, Z., J.J. Kotzin, B. Ramdas, S. Chen, S. Nelanuthala, L.R. Palam, R. Pandey, R.S. Mali, Y. Liu, M.R. Kelley, G. Sandusky, M. Mohseni, A. Williams, J. Henao-Mejia, and R. Kapur. 2018. Inhibition of Inflammatory Signaling in Tet2 Mutant Preleukemic Cells Mitigates Stress-Induced Abnormalities and Clonal Hematopoiesis. Cell Stem Cell 23:833–849 e835.

Cancer Genome Atlas Research, N., T.J. Ley, C. Miller, L. Ding, B.J. Raphael, A.J. Mungall, A. Robertson, K. Hoadley, T.J. Triche, Jr., P.W. Laird, J.D. Baty, L.L. Fulton, R. Fulton, S.E. Heath, J. Kalicki-Veizer, C. Kandoth, J.M. Klco, D.C. Koboldt, K.L. Kanchi, S. Kulkarni, T.L. Lamprecht, D.E. Larson, L. Lin, C. Lu, M.D. McLellan, J.F. McMichael, J. Payton, H. Schmidt, D.H. Spencer, M.H. Tomasson, J.W. Wallis, L.D. Wartman, M.A. Watson, J. Welch, M.C. Wendl, A. Ally, M. Balasundaram, I. Birol, Y. Butterfield, R. Chiu, A. Chu, E. Chuah, H.J. Chun, R. Corbett, N. Dhalla, R. Guin, A. He, C. Hirst, M. Hirst, R.A. Holt, S. Jones, A. Karsan, D. Lee, H.I. Li, M.A. Marra, M. Mayo, R.A. Moore, K. Mungall, J. Parker, E. Pleasance, P. Plettner, J. Schein, D. Stoll, L. Swanson, A. Tam, N. Thiessen, R. Varhol, N. Wye, Y. Zhao, S. Gabriel, G. Getz, C. Sougnez, L. Zou, M.D. Leiserson, F. Vandin, H.T. Wu, F. Applebaum, S.B. Baylin, R. Akbani, B.M. Broom, K. Chen, T.C. Motter, K. Nguyen, J.N. Weinstein, N. Zhang, M.L. Ferguson, C. Adams, A. Black, J. Bowen, J. Gastier-Foster, T. Grossman, T. Lichtenberg, L. Wise, T. Davidsen, J.A. Demchok, K.R. Shaw, M. Sheth, H.J. Sofia, L. Yang, J.R. Downing, and G. Eley. 2013. Genomic and epigenomic landscapes of adult de novo acute myeloid leukemia. N Engl J Med 368:2059–2074.

Carey, A., D.K.t. Edwards, C.A. Eide, L. Newell, E. Traer, B.C. Medeiros, D.A. Pollyea, M.W. Deininger, R.H. Collins, J.W. Tyner, B.J. Druker, G.C. Bagby, S.K. McWeeney, and A. Agarwal. 2017. Identification of Interleukin-1 by Functional Screening as a Key Mediator of Cellular Expansion and Disease Progression in Acute Myeloid Leukemia. Cell Rep 18:3204–3218.

Chambers, S.M., N.C. Boles, K.Y. Lin, M.P. Tierney, T.V. Bowman, S.B. Bradfute, A.J. Chen, A.A. Merchant, O. Sirin, D.C. Weksberg, M.G. Merchant, C.J. Fisk, C.A. Shaw, and M.A. Goodell. 2007. Hematopoietic fingerprints: an expression database of stem cells and their progeny. Cell Stem Cell 1:578–591.

Corces, M.R., H.Y. Chang, and R. Majeti. 2017. Preleukemic Hematopoietic Stem Cells in Human Acute Myeloid Leukemia. Front Oncol 7:263.

Dobin, A., C.A. Davis, F. Schlesinger, J. Drenkow, C. Zaleski, S. Jha, P. Batut, M. Chaisson, and T.R. Gingeras. 2012. STAR: ultrafast universal RNA-seq aligner. Bioinformatics 29:15–21.

Ernst, T., A. Chase, K. Zoi, K. Waghorn, C. Hidalgo-Curtis, J. Score, A. Jones, F. Grand, A. Reiter, A. Hochhaus, and N.C. Cross. 2010. Transcription factor mutations in myelodysplastic/myeloproliferative neoplasms. Haematologica 95:1473–1480.

Fleenor, C.J., K. Higa, M.M. Weil, and J. DeGregori. 2015a. Evolved Cellular Mechanisms to Respond to Genotoxic Insults: Implications for Radiation-Induced Hematologic Malignancies. Radiat Res 184:341–351.

Fleenor, C.J., A.I. Rozhok, V. Zaberezhnyy, D. Mathew, J. Kim, A.C. Tan, I.D. Bernstein, and J. DeGregori. 2015b. Contrasting roles for C/EBPalpha and Notch in irradiation-induced multipotent hematopoietic progenitor cell defects. Stem Cells 33:1345–1358.

Fleischman, A.G., K.J. Aichberger, S.B. Luty, T.G. Bumm, C.L. Petersen, S. Doratotaj, K.B. Vasudevan, D.H. LaTocha, F. Yang, R.D. Press, M.M. Loriaux, H.L. Pahl, R.T. Silver, A. Agarwal, T. O’Hare, B.J. Druker, G.C. Bagby, and M.W. Deininger. 2011. TNFalpha facilitates clonal expansion of JAK2V617F positive cells in myeloproliferative neoplasms. Blood 118:6392–6398.

Gan, B., and R.A. DePinho. 2009. mTORC1 signaling governs hematopoietic stem cell quiescence. Cell Cycle 8:1003–1006.

Garcia-Prat, L., P. Sousa-Victor, and P. Munoz-Canoves. 2017. Proteostatic and Metabolic Control of Stemness. Cell Stem Cell 20:593–608.

Giladi, A., F. Paul, Y. Herzog, Y. Lubling, A. Weiner, I. Yofe, D. Jaitin, N. Cabezas-Wallscheid, R. Dress, F. Ginhoux, A. Trumpp, A. Tanay, and I. Amit. 2018. Single-cell characterization of haematopoietic progenitors and their trajectories in homeostasis and perturbed haematopoiesis. Nat Cell Biol 20:836–846.

Gothert, J.R., S.E. Gustin, M.A. Hall, A.R. Green, B. Gottgens, D.J. Izon, and C.G. Begley. 2005. In vivo fate-tracing studies using the Scl stem cell enhancer: embryonic hematopoietic stem cells significantly contribute to adult hematopoiesis. Blood 105:2724–2732.

Gu, X., Q. Ebrahem, R.Z. Mahfouz, M. Hasipek, F. Enane, T. Radivoyevitch, N. Rapin, B. Przychodzen, Z. Hu, R. Balusu, C.V. Cotta, D. Wald, C. Argueta, Y. Landesman, M.P. Martelli, B. Falini, H. Carraway, B.T. Porse, J. Maciejewski, B.K. Jha, and Y. Saunthararajah. 2018. Leukemogenic nucleophosmin mutation disrupts the transcription factor hub that regulates granulomonocytic fates. J Clin Invest 128:4260–4279.

Gu, Z., R. Eils, and M. Schlesner. 2016. Complex heatmaps reveal patterns and correlations in multidimensional genomic data. Bioinformatics 32:2847–2849.

Guo, H., O. Ma, N.A. Speck, and A.D. Friedman. 2012. Runx1 deletion or dominant inhibition reduces Cebpa transcription via conserved promoter and distal enhancer sites to favor monopoiesis over granulopoiesis. Blood 119:4408–4418.

Hasemann, M.S., F.K. Lauridsen, J. Waage, J.S. Jakobsen, A.K. Frank, M.B. Schuster, N. Rapin, F.O. Bagger, P.S. Hoppe, T. Schroeder, and B.T. Porse. 2014. C/EBPalpha is required for long-term self-renewal and lineage priming of hematopoietic stem cells and for the maintenance of epigenetic configurations in multipotent progenitors. PLoS Genet 10:e1004079.

Henry, C.J., M. Casas-Selves, J. Kim, V. Zaberezhnyy, L. Aghili, A.E. Daniel, L. Jimenez, T. Azam, E.N. McNamee, E.T. Clambey, J. Klawitter, N.J. Serkova, A.C. Tan, C.A. Dinarello, and J. DeGregori. 2015. Aging-associated inflammation promotes selection for adaptive oncogenic events in B cell progenitors. J Clin Invest 125:4666–4680.

Henry, C.J., A. Marusyk, V. Zaberezhnyy, B. Adane, and J. DeGregori. 2010. Declining lymphoid progenitor fitness promotes aging-associated leukemogenesis. Proc Natl Acad Sci U S A 107:21713–21718.

Herault, A., M. Binnewies, S. Leong, F.J. Calero-Nieto, S.Y. Zhang, Y.A. Kang, X. Wang, E.M. Pietras, S.H. Chu, K. Barry-Holson, S. Armstrong, B. Gottgens, and E. Passegue. 2017. Myeloid progenitor cluster formation drives emergency and leukaemic myelopoiesis. Nature 544:53–58.

Hsu, J.I., T. Dayaram, A. Tovy, E. De Braekeleer, M. Jeong, F. Wang, J. Zhang, T.P. Heffernan, S. Gera, J.J. Kovacs, J.R. Marszalek, C. Bristow, Y. Yan, G. Garcia-Manero, H. Kantarjian, G. Vassiliou, P.A. Futreal, L.A. Donehower, K. Takahashi, and M.A. Goodell. 2018. PPM1D Mutations Drive Clonal Hematopoiesis in Response to Cytotoxic Chemotherapy. Cell Stem Cell 23:700–713 e706.

Ito, K., and T. Suda. 2014. Metabolic requirements for the maintenance of self-renewing stem cells. Nat Rev Mol Cell Biol 15:243–256.

Iwama, A., H. Oguro, M. Negishi, Y. Kato, Y. Morita, H. Tsukui, H. Ema, T. Kamijo, Y. Katoh-Fukui, H. Koseki, M. van Lohuizen, and H. Nakauchi. 2004. Enhanced self-renewal of hematopoietic stem cells mediated by the polycomb gene product Bmi-1. Immunity 21:843–851.

Jalbert, E., and E.M. Pietras. 2018. Analysis of Murine Hematopoietic Stem Cell Proliferation During Inflammation. In Cellular Quiescence: Methods and Protocols. H.D. Lacorazza, editor Springer New York, New York, NY. 183–200.

Johnson, P.F. 2005. Molecular stop signs: regulation of cell-cycle arrest by C/EBP transcription factors. J Cell Sci 118:2545–2555.

Kahn, J.D., P.G. Miller, A.J. Silver, R.S. Sellar, S. Bhatt, C. Gibson, M. McConkey, D. Adams, B. Mar, P. Mertins, S. Fereshetian, K. Krug, H. Zhu, A. Letai, S.A. Carr, J. Doench, S. Jaiswal, and B.L. Ebert. 2018. PPM1D-truncating mutations confer resistance to chemotherapy and sensitivity to PPM1D inhibition in hematopoietic cells. Blood 132:1095–1105.

Kang, Y.A., E.M. Pietras, and E. Passegue. 2020. Deregulated Notch and Wnt signaling activates early-stage myeloid regeneration pathways in leukemia. J Exp Med 217:

Kelly, L.M., and D.G. Gilliland. 2002. Genetics of myeloid leukemias. Annu Rev Genomics Hum Genet 3:179–198.

Kirstetter, P., M.B. Schuster, O. Bereshchenko, S. Moore, H. Dvinge, E. Kurz, K. Theilgaard-Monch, R. Mansson, T.A. Pedersen, T. Pabst, E. Schrock, B.T. Porse, S.E. Jacobsen, P. Bertone, D.G. Tenen, and C. Nerlov. 2008. Modeling of C/EBPalpha mutant acute myeloid leukemia reveals a common expression signature of committed myeloid leukemia-initiating cells. Cancer Cell 13:299–310.

Laurenti, E., B. Varnum-Finney, A. Wilson, I. Ferrero, W.E. Blanco-Bose, A. Ehninger, P.S. Knoepfler, P.F. Cheng, H.R. MacDonald, R.N. Eisenman, I.D. Bernstein, and A. Trumpp. 2008. Hematopoietic stem cell function and survival depend on c-Myc and N-Myc activity. Cell Stem Cell 3:611–624.

Liberzon, A., A. Subramanian, R. Pinchback, H. Thorvaldsdóttir, P. Tamayo, and J.P. Mesirov. 2011. Molecular signatures database (MSigDB) 3.0. Bioinformatics 27:1739–1740.

Mantovani, A., C.A. Dinarello, M. Molgora, and C. Garlanda. 2019. Interleukin-1 and Related Cytokines in the Regulation of Inflammation and Immunity. Immunity 50:778–795.

Manz, M.G., and S. Boettcher. 2014. Emergency granulopoiesis. Nat Rev Immunol 14:302–314.

Marusyk, A., M. Casas-Selves, C.J. Henry, V. Zaberezhnyy, J. Klawitter, U. Christians, and J. DeGregori. 2009. Irradiation alters selection for oncogenic mutations in hematopoietic progenitors. Cancer Res 69:7262–7269.

Marusyk, A., C.C. Porter, V. Zaberezhnyy, and J. DeGregori. 2010. Irradiation selects for p53-deficient hematopoietic progenitors. PLoS Biol 8:e1000324.

Mejia-Ramirez, E., and M.C. Florian. 2020. Understanding intrinsic hematopoietic stem cell aging. Haematologica 105:22–37.

Miyamoto, K., K.Y. Araki, K. Naka, F. Arai, K. Takubo, S. Yamazaki, S. Matsuoka, T. Miyamoto, K. Ito, M. Ohmura, C. Chen, K. Hosokawa, H. Nakauchi, K. Nakayama, K.I. Nakayama, M. Harada, N. Motoyama, T. Suda, and A. Hirao. 2007. Foxo3a is essential for maintenance of the hematopoietic stem cell pool. Cell Stem Cell 1:101–112.

Nitta, E., N. Itokawa, S. Yabata, S. Koide, L.B. Hou, M. Oshima, K. Aoyama, A. Saraya, and A. Iwama. 2020. Bmi1 counteracts hematopoietic stem cell aging by repressing target genes and enforcing the stem cell gene signature. Biochem Biophys Res Commun 521:612–619.

Pabst, T., B.U. Mueller, N. Harakawa, C. Schoch, T. Haferlach, G. Behre, W. Hiddemann, D.-E. Zhang, and D.G. Tenen. 2001a. AML1–ETO downregulates the granulocytic differentiation factor C/EBPα in t(8;21) myeloid leukemia. Nature Medicine 7:444–451.

Pabst, T., B.U. Mueller, P. Zhang, H.S. Radomska, S. Narravula, S. Schnittger, G. Behre, W. Hiddemann, and D.G. Tenen. 2001b. Dominant-negative mutations of CEBPA, encoding CCAAT/enhancer binding protein-α (C/EBPα), in acute myeloid leukemia. Nature Genetics 27:263–270.

Peng, H., A. Kasada, M. Ueno, T. Hoshii, Y. Tadokoro, N. Nomura, C. Ito, Y. Takase, H.T. Vu, M. Kobayashi, B. Xiao, P.F. Worley, and A. Hirao. 2018. Distinct roles of Rheb and Raptor in activating mTOR complex 1 for the self-renewal of hematopoietic stem cells. Biochem Biophys Res Commun 495:1129–1135.

Perrotti, D., V. Cesi, R. Trotta, C. Guerzoni, G. Santilli, K. Campbell, A. Iervolino, F. Condorelli, C. Gambacorti-Passerini, M.A. Caligiuri, and B. Calabretta. 2002. BCR-ABL suppresses C/EBPalpha expression through inhibitory action of hnRNP E2. Nat Genet 30:48–58.

Pietras, E.M. 2017. Inflammation: a key regulator of hematopoietic stem cell fate in health and disease. Blood 130:1693–1698.

Pietras, E.M., C. Mirantes-Barbeito, S. Fong, D. Loeffler, L.V. Kovtonyuk, S. Zhang, R. Lakshminarasimhan, C.P. Chin, J.M. Techner, B. Will, C. Nerlov, U. Steidl, M.G. Manz, T. Schroeder, and E. Passegue. 2016. Chronic interleukin-1 exposure drives haematopoietic stem cells towards precocious myeloid differentiation at the expense of self-renewal. Nat Cell Biol 18:607–618.

Pietras, E.M., D. Reynaud, Y.A. Kang, D. Carlin, F.J. Calero-Nieto, A.D. Leavitt, J.M. Stuart, B. Gottgens, and E. Passegue. 2015. Functionally Distinct Subsets of Lineage-Biased Multipotent Progenitors Control Blood Production in Normal and Regenerative Conditions. Cell Stem Cell 17:35–46.

Porse, B.T., D. Bryder, K. Theilgaard-Monch, M.S. Hasemann, K. Anderson, I. Damgaard, S.E. Jacobsen, and C. Nerlov. 2005. Loss of C/EBP alpha cell cycle control increases myeloid progenitor proliferation and transforms the neutrophil granulocyte lineage. J Exp Med 202:85–96.

Pundhir, S., F.K. Bratt Lauridsen, M.B. Schuster, J.S. Jakobsen, Y. Ge, E.M. Schoof, N. Rapin, J. Waage, M.S. Hasemann, and B.T. Porse. 2018. Enhancer and Transcription Factor Dynamics during Myeloid Differentiation Reveal an Early Differentiation Block in Cebpa null Progenitors. Cell Rep 23:2744–2757.

Ritchie, M.E., B. Phipson, D. Wu, Y. Hu, C.W. Law, W. Shi, and G.K. Smyth. 2015. limma powers differential expression analyses for RNA-sequencing and microarray studies. Nucleic Acids Research 43:e47–e47.

Sawen, P., S. Lang, P. Mandal, D.J. Rossi, S. Soneji, and D. Bryder. 2016. Mitotic History Reveals Distinct Stem Cell Populations and Their Contributions to Hematopoiesis. Cell Rep 14:2809–2818.

Sergushichev, A.A. 2016. An algorithm for fast preranked gene set enrichment analysis using cumulative statistic calculation. 060012.

Signer, R.A., J.A. Magee, A. Salic, and S.J. Morrison. 2014. Haematopoietic stem cells require a highly regulated protein synthesis rate. Nature 509:49–54.

Srinivas, S., T. Watanabe, C.-S. Lin, C.M. William, Y. Tanabe, T.M. Jessell, and F. Costantini. 2001. Cre reporter strains produced by targeted insertion of EYFP and ECFP into the ROSA26 locus. BMC Developmental Biology 1:4.

Sun, J., A. Ramos, B. Chapman, J.B. Johnnidis, L. Le, Y.J. Ho, A. Klein, O. Hofmann, and F.D. Camargo. 2014. Clonal dynamics of native haematopoiesis. Nature 514:322–327.

Varnum-Finney, B., C. Brashem-Stein, and I.D. Bernstein. 2003. Combined effects of Notch signaling and cytokines induce a multiple log increase in precursors with lymphoid and myeloid reconstituting ability. Blood 101:1784–1789.

Vas, V., K. Senger, K. Dorr, A. Niebel, and H. Geiger. 2012. Aging of the microenvironment influences clonality in hematopoiesis. PLoS One 7:e42080.

Weber, A., P. Wasiliew, and M. Kracht. 2010. Interleukin-1 (IL-1) Pathway. 3:cm1-cm1.

Ye, M., H. Zhang, G. Amabile, H. Yang, P.B. Staber, P. Zhang, E. Levantini, M. Alberich-Jordà, J. Zhang, A. Kawasaki, and D.G. Tenen. 2013. C/EBPa controls acquisition and maintenance of adult haematopoietic stem cell quiescence. Nature Cell Biology 15:385–394.

Yoshihara, H., F. Arai, K. Hosokawa, T. Hagiwara, K. Takubo, Y. Nakamura, Y. Gomei, H. Iwasaki, S. Matsuoka, K. Miyamoto, H. Miyazaki, T. Takahashi, and T. Suda. 2007. Thrombopoietin/MPL signaling regulates hematopoietic stem cell quiescence and interaction with the osteoblastic niche. Cell Stem Cell 1:685–697.

Yu, G., L.-G. Wang, Y. Han, and Q.-Y. He. 2012. clusterProfiler: an R package for comparing biological themes among gene clusters. Omics : a journal of integrative biology 16:284–287.

Zhang, D.-E., P. Zhang, N.-d. Wang, C.J. Hetherington, G.J. Darlington, and D.G. Tenen. 1997. Absence of granulocyte colony-stimulating factor signaling and neutrophil development in CCAAT enhancer binding protein α-deficient mice. Proceedings of the National Academy of Sciences 94:569–574.

Zhang, M., F. Liu, P. Zhou, Q. Wang, C. Xu, Y. Li, L. Bian, Y. Liu, J. Zhou, F. Wang, Y. Yao, Y. Fang, and D. Li. 2019. The MTOR signaling pathway regulates macrophage differentiation from mouse myeloid progenitors by inhibiting autophagy. Autophagy 15:1150–1162.

Zhang, P., J. Iwasaki-Arai, H. Iwasaki, M.L. Fenyus, T. Dayaram, B.M. Owens, H. Shigematsu, E. Levantini, C.S. Huettner, J.A. Lekstrom-Himes, K. Akashi, and D.G. Tenen. 2004. Enhancement of hematopoietic stem cell repopulating capacity and self-renewal in the absence of the transcription factor C/EBP alpha. Immunity 21:853–863.

Zheng, R., A.D. Friedman, M. Levis, L. Li, E.G. Weir, and D. Small. 2004. Internal tandem duplication mutation of FLT3 blocks myeloid differentiation through suppression of C/EBPα expression. Blood 103:1883–1890.

